# Test-retest reliability of TMS motor evoked responses and silent periods during explosive voluntary isometric contractions

**DOI:** 10.1101/2024.08.12.607577

**Authors:** F. Castelli, O.S. Mian, A. Bruton, A.C. Chembila Valappil, N.A. Tillin

## Abstract

**Purpose:** This study assessed the intraclass correlation coefficient (ICC) and coefficient of variation (CV) test-retest reliability of TMS responses MEPs and silent periods at early, middle, and late phases of the rising time-torque curve during explosive voluntary contractions. We also investigated how the number of consecutively averaged measurements over 3 to 15 separate contractions influenced reliability.

**Methods:** On two separate occasions 3-7 days apart, 14 adults completed 48 isometric explosives (1-s) contractions of the knee extensors, superimposed with TMS. The TMS elicited an MEP and silent period in the quadriceps muscles at 45 (early), 115 (middle), or 190 ms (late) during each contraction. TMS was also superimposed at the plateau of 15 separate MVCs. Test-retest ICC and CV were calculated for consecutively averaged MEPs and silent periods.

**Results:** No one condition was more reliable than another. For MEP amplitude, in all conditions except the explosive late phase, ICCs generally increased, and CV decreased, with an increase in the number of averaged contractions, and were >0.50 ICC and <15% CV within 7 contractions. For silent period, ICCs and CVs were unaffected by the number of consecutively averaged contractions and remained >0.50 ICC and <10% CV.

**Conclusion:** Test-retest reliability of TMS responses is comparable between phases of explosive contraction and at the plateau of MVC. To enhance MEP reliability, with the increases he number of averaged contractions of except the explosive late phase, we recommend studies average data across > the 3-5 contractions typically reported in the literature for studies using TMS during MVCs.

## Introduction

The rate of torque development (RTD) measures the ability of muscles to rapidly increase force around a joint (Tillin and Folland, 2014), and is functionally important where time to develop force is limited, such as sprinting (Tillin, Pain and Folland, 2013) or balance recovery (Sundstrup *et al*., 2010). RTD is often measured in early (0-50 ms), middle (50-100 ms) and late (>100 ms) phases of explosive contraction performed from rest (Maffiuletti *et al*., 2016), and different physiological factors limit the RTD in these separate phases. One limiting factor is the neural drive to muscle, which positively correlates with early- and middle-phase RTD (Folland, Buckthorpe and Hannah, 2014; Del Vecchio *et al*., 2019). Despite the relevance of neural drive to RTD, the corticospinal mechanisms affecting RTD are not well established.

Transcranial magnetic stimulation (TMS), superimposed during voluntary contractions, elicits a motor-evoked potential (MEP) in the surface EMG signal of the target contracting muscle. The MEP amplitude is thought to reflect corticospinal excitability (Rossini *et al*., 2015), whilst a period of electrical inactivity immediately after the MEP, referred to as the silent period (Damron *et al*., 2008), is thought to reflect corticospinal inhibitory mechanisms (Säisänen *et al*., 2008). Despite extensive use of TMS to explore corticospinal excitability/inhibition at the plateau of a maximal voluntary contraction (MVC; Todd, Taylor and Gandevia, 2016), TMS has not been commonly used during the rising torque of explosive voluntary contractions, where RTD is typically measured.

Before using TMS to assess corticospinal excitability/inhibition during explosive contractions, it is important to establish the test-retest absolute and relative reliability of TMS responses (MEP amplitude and silent period) during such conditions; however, this has not been done. In contrast, the test-retest reliability for TMS responses at the plateau of MVCs has been investigated in various muscles. For absolute and normalised MEP amplitude at the MVC plateau in the lower limbs, studies have reported moderate inter-class correlation (ICC; 0.52-0.79) (Mileva, Sumners and Bowtell, 2012; Souron *et al*., 2016) and coefficient of variation (CV) of 10-11% (Souron *et al*., 2016). For the silent period at the MVC plateau, the test-retest ICC has ranged from moderate in the vastus lateralis (0.61-0.70; (Di Virgilio *et al*., 2022)) to excellent in the soleus and tibialis anterior (0.93-0.95; Mileva, Sumners and Bowtell, 2012; Souron *et al*., 2016) and one study has reported a CV of 8.6% (Souron *et al*., 2016) in the tibialis anterior. It is unclear whether the reliability of MEP amplitudes and silent periods during explosive contractions will be comparable to that observed at the MVC plateau. Typically, the reliability of torque and EMG amplitude measurements is lower in early compared to later phases of explosive contraction, and generally lower for explosive contractions compared to the MVC plateau (De Ruiter *et al*., 2004; Buckthorpe *et al*., 2012), so a similar pattern may be observed for the reliability of MEP amplitudes and silent periods.

TMS responses are typically averaged across multiple separate stimuli (in separate contractions) to minimise the influence of random variation and improve reliability. Thus, when assessing the test-retest reliability of MEP amplitudes and silent periods, it is important to consider how many stimulations might be required to achieve adequately reliable averages. A relatively high number of stimuli (≥20) are needed to obtain moderate to good test-retest reliability (ICC = 0.50 - 0.75; Goldsworthy, Hordacre and Ridding, 2016; Biabani *et al*., 2018) of MEP amplitudes recorded when the participant is passive. In contrast, as few as four stimuli have been shown to produce good (≥0.75) test-retest ICC for MEP amplitude during submaximal contractions held at a constant force (Wheaton *et al*., 2009; Lewis, Signal and Taylor, 2013; Temesi, Ly and Millet, 2017). The effect of the number of contractions on test-retest reliability of average TMS responses has not been determined during either MVC or explosive contractions. Previous studies of TMS responses during MVCs have typically only collected and averaged 3-5 responses (Luc *et al*., 2014; Tallent *et al*., 2017; Škarabot *et al*., 2019). This could be due to time constraints or concerns over fatigue with multiple contractions; however, it may be at the expense of poor reliability. Thus, there is a need to assess the effect of the number of contractions on the test-retest reliability of average TMS responses during MVCs and explosive contractions.

In studies of TMS responses, measures are typically averaged across all stimulations obtained within a testing session. However, a subset of contractions are typically used for studies of explosive contractions, with the best three out of ten contractions being recommended for averaging RTD measurements (Maffiuletti *et al*., 2016). Therefore, for studies of MEPs recorded during explosive contractions, it may be desirable to investigate MEPs obtained from the best three contractions rather than from all contractions, and investigation into the reliability of this approach is necessary.

This study aims to assess the absolute (ICC) and relative (CV) test-retest reliability of TMS responses (MEP amplitude and silent period) recorded at different time points (early, middle, and late phases) during explosive voluntary contractions and at the plateau of MVCs. As part of this aim, we intend to (i) document how the number of consecutive contractions (between 3-15) over which TMS responses are averaged, influences test-retest reliability; and (ii) document test-retest reliability of TMS responses averaged across the best 3, out of 10, contractions. We determine the best three as those with the highest torque (Torque averaging method) or EMG RMS (EMG averaging method) prior to MEP on-set.

## Methods

### Participants

Fourteen participants, nine males (age 31 ± 5 years, height, 178.7 ± 6.6cm, and mass 79.2 ± 4.7kg) and five females (age 29 ± 6 years, height, 165.5 ± 5.5cm, and mass 59.3 ± 6.2kg) were recruited to take part in this study. All participants habitually performed 120-180 minutes of moderate to high-intensity activity per week, and were deemed recreationally active (McKay *et al*., 2022). Participants were also free from injury and disease (screened via a questionnaire adapted from Balady *et al*., 1998) and free from contraindications to TMS (screened via a questionnaire adapted from Rossi *et al*., 2011). Due to changes in endogenous hormones throughout the menstrual cycle potentially affecting neuromuscular responses to TMS (Ansdell *et al*., 2019), female participants undertook experimental trials exclusively during their self-reported early to mid-follicular phase (first ten days from the first day of menstruation), when endogenous hormone concentration is low and relatively stable (de Jonge, Thompson and Han, 2019). The University of Roehampton ethics committee approved the study, and all participants provided written informed consent before participating.

### Overview

Participants visited the laboratory on three separate occasions and were asked to avoid strenuous exercise and alcohol consumption for 24 hours before each visit. Each session lasted approximately 120-150 minutes, with consecutive sessions separated by 3-7 days. The first visit was a familiarisation session, and the second and third were measurement sessions. The measurement sessions were performed at a consistent time of day and involved an identical protocol, with measurements used to assess the test-retest reliability of the variables of interest. Each session involved knee extensor torque and EMG measurement during maximal voluntary and explosive contractions, using TMS to obtain superimposed MEP and femoral nerve stimulation to obtain compound muscle action potentials at rest.

### Torque measurements and surface electromyography (EMG)

Participants were tightly secured in a custom-built strength testing chair (Fig. 6b in Maffiuletti *et al*., 2016) with a waist belt and shoulder straps. The hip and knee angles were set at 100° and 105°, respectively (full extension being 180°). All contractions were isometric knee extensions performed with the right leg. An ankle strap joined to a calibrated S-shaped load cell (FSB-1.5kN, Force Logic, Reading, UK) was secured 4 cm proximal to the medial malleolus. The force signal was amplified (x375) and then sampled at 2000 Hz (Mirco3 1401 and Spike2 v.8; CED., Cambridge, UK). Offline, the force was filtered (fourth-order low-pass Butterworth, 250 Hz cut-off), corrected for limb weight, and multiplied by the external moment arm to calculate joint torque.

The skin was prepared by shaving, cleaning (70% ethanol), and lightly abrading the area where EMG electrodes were placed. A single, bipolar silver-silver-chloride gel-electrode configuration (2-cm diameter and 2-cm inter-electrode distance; Dual Electrode, Noraxon, Arizona, USA) was placed over the belly of each of the rectus femoris (RF), vastus lateralis (VL) and vastus medialis (VM), based on SENIAM guidelines (Hermens *et al*., 1999). The EMG system has an inherent 312-ms delay, so whilst suitable for measurements involved in the study, EMG signals sampled by it could not be used to detect activation on-set and trigger the TMS in real-time during the explosive contractions (explained below in Experimental procedures). Thus, a second wired bipolar EMG electrode (2 cm diameter and 2 cm inter-electrode distance; Dual Electrode, Biometrics Ltd, Gwent, UK) was placed on the belly of the VM to trigger the TMS during contraction. Electrode locations were marked with a permanent marker pen, and participants were asked to maintain these marks throughout the study by re-applying them if necessary. The three wireless EMG signals were transmitted to a desktop receiver (TeleMYO D.T.S., Noraxon, Arizona, USA) and sampled, along with the single wired EMG signal, at 2000 Hz via the same A/D convertor and software as the force signal. Once offline, wireless EMG signals were filtered (fourth-order Butterworth, band-pass, 6-500 Hz) and time-corrected for the 312-ms delay inherent in the Noraxon system.

### Transcranial magnetic stimulation (TMS)

TMS with a 1-ms pulse width was delivered via a double cone coil (110-mm Magstim 200, Whitland, UK) over the scalp in an optimal position to elicit MEPs in the right quadriceps muscles. The following procedures were completed in every session. Participants wore a swim cap, and the vertex of the head – identified as 50% of the distance between nasion and inion and between right and left perpendicular point – was marked on the swim cap. A 5-by-5-cm grid with 1-cm spacing between grid lines was drawn on the swim cap, lateral (left hemisphere) and posterior from the vertex. The coil was moved posteriorly and laterally from the vertex in ∼0.5-cm steps, and in each position, the participant completed four submaximal voluntary contractions at 20% MVC torque (established in the warm-up; see below) with superimposed TMS on each contraction at a submaximal (range 50-60%) stimulator output. The position determined the optimal coil position for the rest of the session, which was subjectively the highest consistent MEP amplitudes (peak-to-peak) over the four superimposed contractions for all three muscles (RF, VL and VM). The optimal coil position was marked by drawing the edge of the coil over the swim cap, ensuring accurate relocation throughout the session. The active motor threshold (AMT) was then determined via a series of 20% MVC torque contractions superimposed with TMS stimulator output starting at 39%. The AMT was defined as the minimum TMS intensity required to elicit five visible MEPs amongst the background EMG activity of the VM and RF, out of ten consecutive superimposed contractions. If the muscles had less or more than five visible MEPs, the machine intensity was reduced or increased by 2% of the machine output. We prioritised the VM and RF (not the VL) as these muscles produced more consistent and visible MEPs in pilot testing. Where it was impossible to match AMT for VM and RF, we settled on being one visible MEP away from 5/10 (e.g., 6/10 VM and 4/10 RF). The same investigator held the coil by hand throughout the measurement sessions, continuously monitoring its position and orientation. For TMS delivered during maximal and explosive contractions (see experimental protocol), the intensity was set at 140% of the stimulator output at AMT (Groppa *et al*., 2012; Rossini *et al*., 2015; Rossi *et al*., 2020). These procedures were repeated for all sessions.

### Femoral nerve electrical stimulation

Single, square-wave pulses (200 µs duration) were delivered (DS7AH, Digitimer, Hertfordshire, UK) over the femoral nerve in the inguinal triangle to evoke twitch contractions and obtain compound muscle action potentials (M-waves) at rest. The anode (5 x 8 cm carbon rubber; EMS Physio Ltd., Oxfordshire, UK) was placed over the head of the greater trochanter. The optimal location of the cathode (1 cm diameter tip; S1 Compex Motor PointPen, Digitimer, UK) was determined as that which evoked the greatest peak twitch torque for a given submaximal stimulation intensity (80-120 mA). The cathode was taped down and held in position by the same investigator as a series of twitches at incremental intensities were evoked until there was a plateau in the peak-to-peak M-wave amplitude (M_max_) of all three muscles (RF, VL, VM). The stimulator intensity was then increased to 150% of the intensity at M_max_, ensuring supramaximal intensity. Three supramaximal twitch contractions were then evoked, each separated by 15 s, and M_max_ was averaged across the three contractions for each muscle. The procedures were repeated for all sessions.

### Experimental protocol

Participants first completed a warm-up involving a series of explosive, submaximal, and maximal voluntary contractions (the latter being used to establish MVC torque), followed by procedures for obtaining optimal TMS coil position, ATM, and M_max_. Participants then completed a series of MVCs and explosive voluntary contractions with and without superimposed TMS. The instruction for MVCs was to "push as hard as possible" for 3-5 s and explosive contractions to "push as fast and hard as possible", emphasising fast for 1 s. The protocol was organised into three blocks of contractions (Fig. 1). Each block involved 24 explosive contractions (8 without and 16 with superimposed TMS) and 8 MVCs (3 without and 5 with superimposed TMS), distributed across four sets. Each set involved six explosive contractions (the final four with TMS stimulation) and two MVCs. In sets 1-3, one of the two MVCs had superimposed TMS (randomly ordered), and in set four, both MVCs had superimposed TMS (Fig. 1). Participants rested 10 s between explosive contractions, 30 s between MVCs, 120 s between sets, and 300 s between blocks. Each block was identical except for the timing of TMS application during explosive contractions, with each block using a different TMS timing condition for these contractions (see next section). The order of the blocks was randomised across participants but held constant across sessions for the same participant. Overall, the 3-block protocol yielded 24 explosive contractions and 9 MVCs without TMS, and 48 explosive contractions (16 per stimulus time condition) and 15 MVCs with TMS.

**Fig 1.**
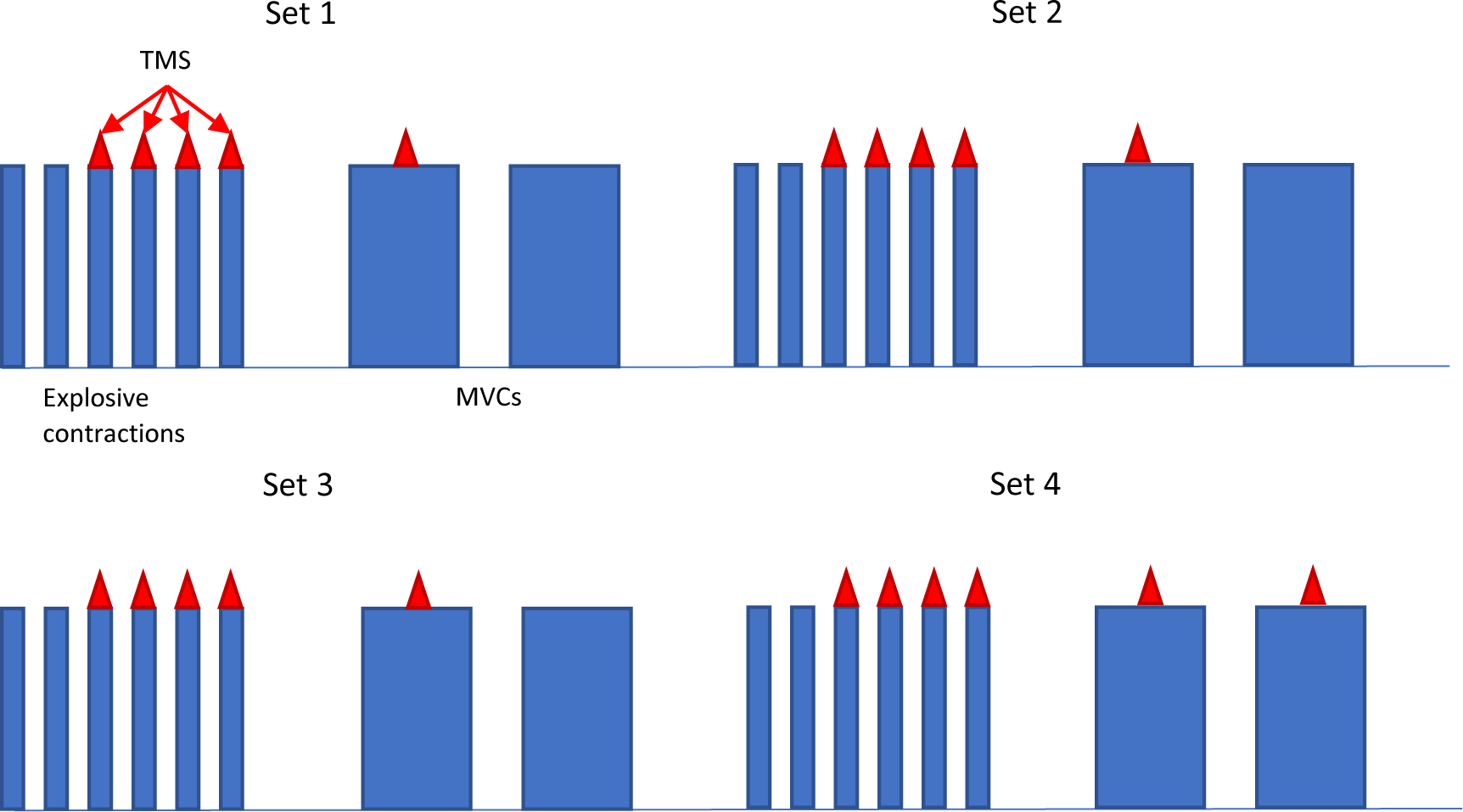
The order of experimental contractions in a block. There were 3 blocks per experimental session and each block involved a different condition for explosive contractions performed with superimposed TMS. In one condition/block TMS occurred in the early phase, in another the middle phase, and in another the late phase, of explosive contraction. The order that blocks/conditions were completed was randomised. Each block consisted of 4 sets of contractions as shown. The thin blue bar represents explosive contractions, while the wide blue bar represents MVCs. The red triangle on the top of the blue bars represents the superimposed TMS

For superimposed MVCs, the TMS was triggered manually by the same experienced investigator during the torque plateau of the MVC. For explosive contractions, the TMS was triggered automatically when the VM EMG signal of the wired system exceeded a set threshold. The threshold was set in the Spike2 software as the lowest amplitude for that session above the highest peaks and troughs of the baseline noise. Commencing from manually detected EMG on-set (0 ms; determined via the methods of Tillin *et al*., 2010), the threshold was exceeded after ∼5 ms, and triggered the TMS at either 8, 78, or 153 ms (accounting for a 3 ms system delay), for the early, middle, and late-phase conditions, respectively (Fig 2). The centre of the evoked MEPs would then occur at ∼45 (early), 115 (middle), or 190 ms (late) from manually detected EMG on-set (Fig. 2).

**Fig 2.**
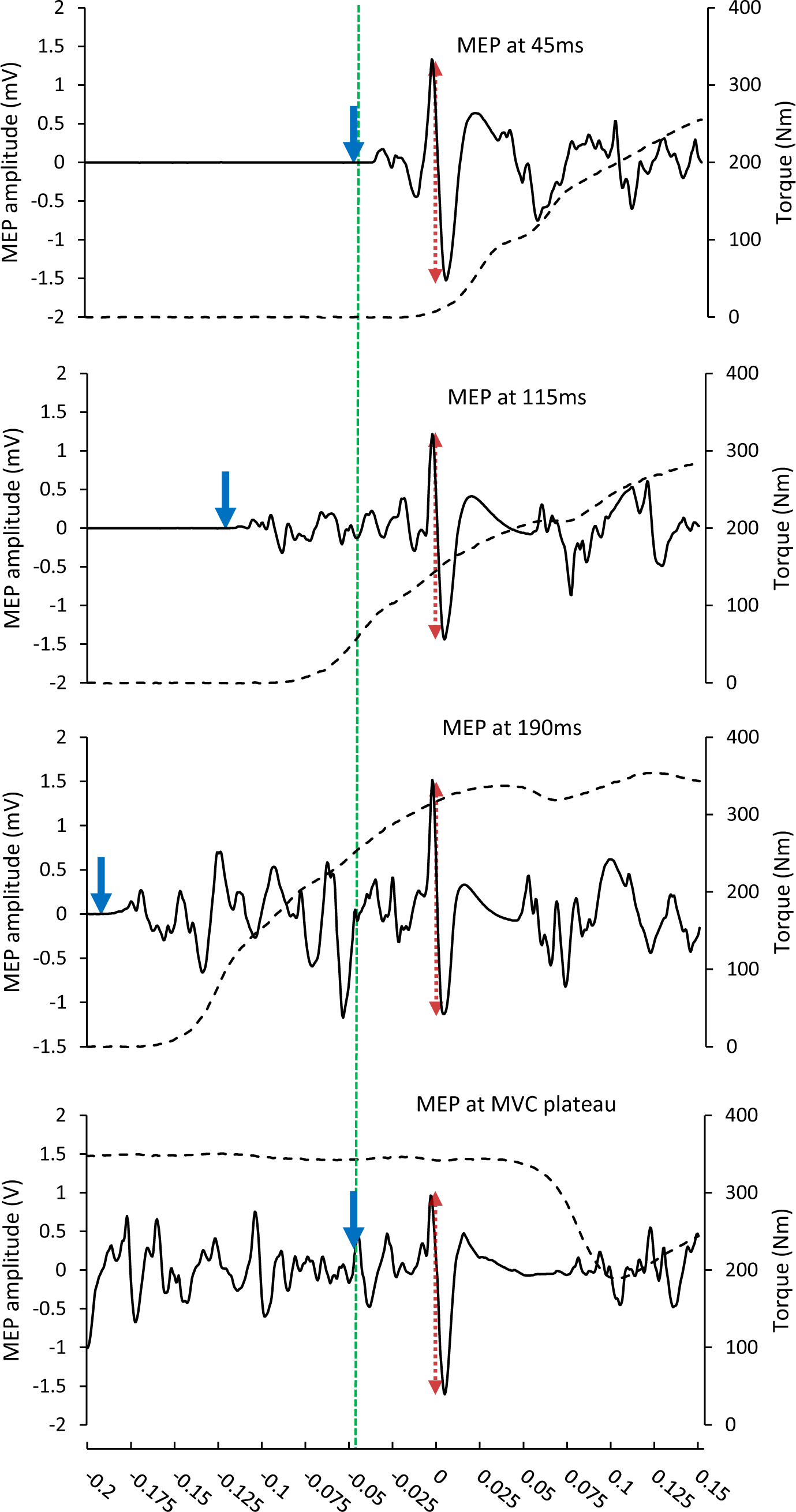
Representative traces of motor evoked potentials (full black line) in the vastus medialis and knee extensor torque (empty dotted line) in each of the four conditions: Early (45 ms), middle (115 ms), and Late (190 ms) from EMG on-set during explosive contractions, and at MVC plateau. The vertical black line across all graphs represents the time of stimulation. The blue line in each graph represents when the TMS was triggered in each condition. The red dotted arrows represent the peak-to-peak amplitude of the MEPs used for analysis

### Data screening and analysis

Out of 15 superimposed MVCs, only those where the TMS was delivered >90% of MVC torque were used for further analysis. Out of the 16 superimposed explosive contractions per condition, only those that met the following criteria were used for further analysis: (i) average baseline force did not change by >2 Nm during the 200 ms preceding manually detected force on-set (detected as in Tillin *et al*., 2010); and (ii) there was a genuine attempt at an explosive contraction. A genuine attempt at an explosive contraction was defined as the instantaneous slope of torque-time, just prior to MEP on-set, being within 3 standard deviations (SD) of mean instantaneous slope at the same time point for contractions without TMS. The time points for measuring instantaneous slope were 30, 105, and 180 ms after manually detected torque on-set, for early-, middle- and late-phase conditions, respectively. Given the criteria above, the maximum number of MEPs analysed further in each condition was limited to the number of usable contractions in the participant with the lowest number of usable contractions. This was 9 (early), 10 (middle), and 15 (late) for the explosive contractions, and 15 for the MVCs.

For useable contractions, MEP amplitude was defined as the peak-to-trough of the superimposed response (Fig. 2) and is reported in absolute terms and normalised to M_max_. The silent period was measured from the point of stimulation to the resumption of EMG activity after the MEP off set. To determine the resumption of EMG activity, the second derivative of EMG amplitude over time was established (1-ms time constant) and then the signal was rectified. EMG activity resumption was defined when the amplitude of the second derivative increased above 5 SD of the mean baseline (calculated in a 500 ms period prior to the contraction) for 70% of the next 10 ms (adapted from Damron *et al*., 2008; Fig.3). This automated process was confirmed via manual inspection of the signal as recommended by Damron et al., (2008). MEP amplitude (absolute and normalised to M_max_ separately) and silent periods were extracted on a muscle level before being averaged across the three muscles to obtain a quadriceps mean. This is standard practice in studies looking to relate general quadriceps EMG data to net knee extensor torque and RTD (Tillin *et al*., 2010; Tillin, Pain and Folland, 2012, 2013, 2018; Folland, Buckthorpe and Hannah, 2014; Behrens *et al*., 2015; Morales-Artacho *et al*., 2018; Cossich and Maffiuletti, 2020).

**Fig 3.**
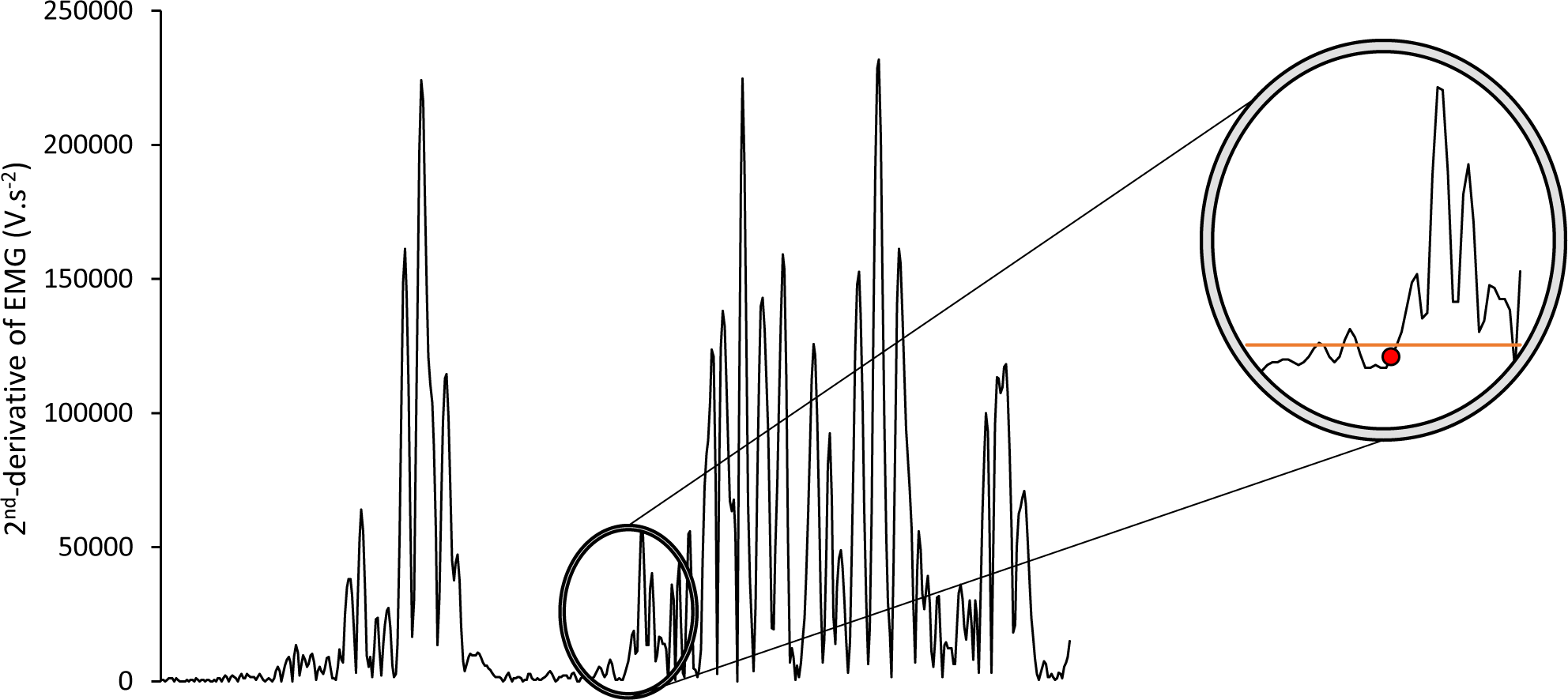
Example of how the silent period off-set was detected in the VM of one participant. The data shows the second derivative of the EMG-time trace after rectification. Circled data are an example of SP off-set detection. The orange line represents the detection threshold (5 SD of the mean baseline), whilst the red dot, which is the detected off-set, represents the last time point before the EMG signal crosses threshold, and remains above the threshold for ≥70% of the data points recorded in the next 10 ms

**Fig 4.**
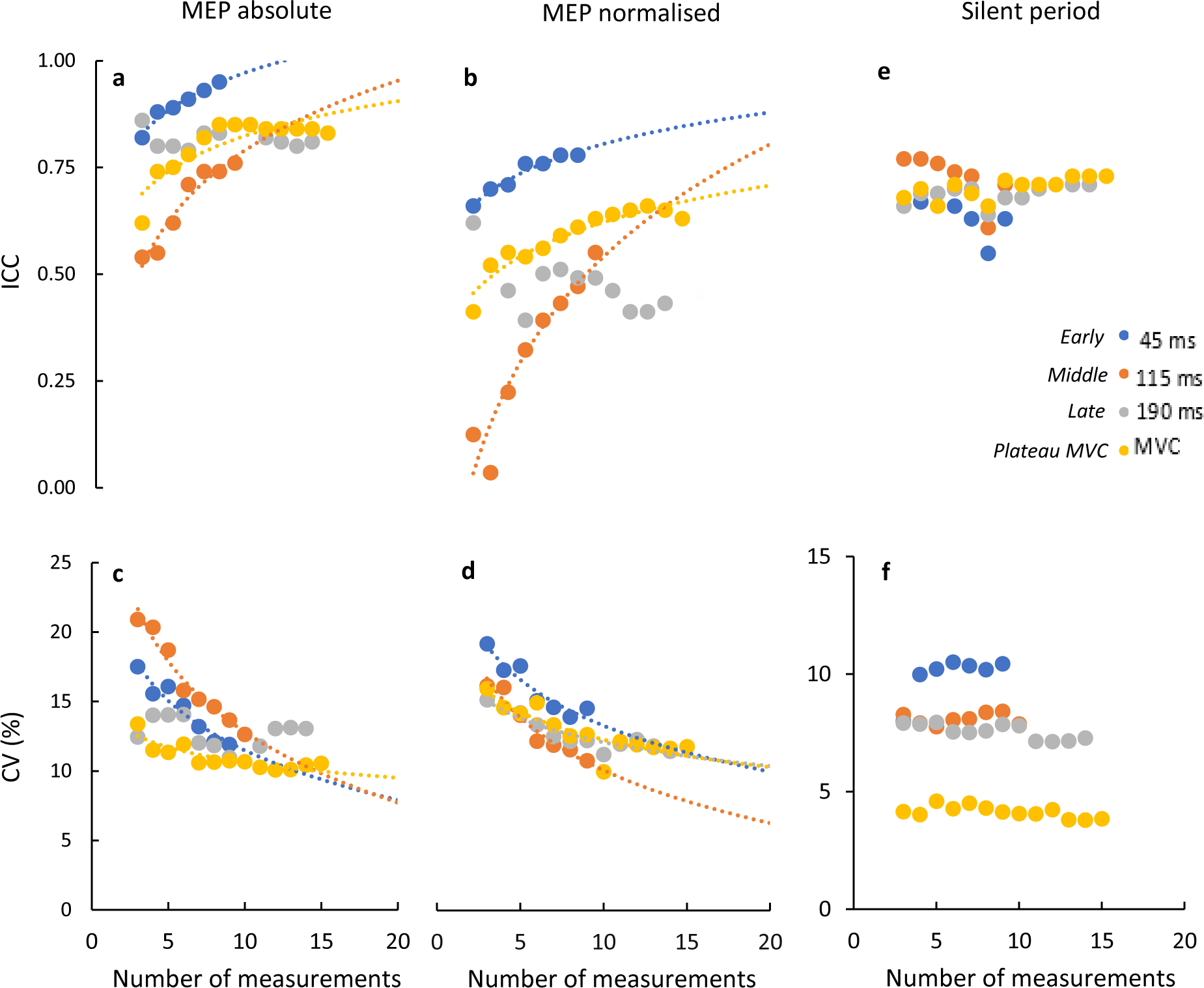
test-retest ICC (A, B, E) and CV (C, D, F) for dependent variables as a function of the number of consecutive contractions over which data are averaged. Dependent variables are absolute (A and C) and normalised (B and D) MEP amplitude, and silent period (E and F). For normalised data, MEP amplitude is normalised to maximal M-wave. MEP amplitudes and silent periods were recorded at the plateau of MVCs, and at 3 different time points from activation on-set during explosive contractions: early (45 ms), middle (115 ms), and Late (190 ms). Logarithmic functions (y=a ln(x)+k) were fitted to data and are plotted as a dotted line for conditions where the fit was significant (p<0.05)

### Averaging methods

For each condition (early, middle, and late explosive phases, and at MVC plateau), mean quadriceps consecutive running averages of the dependent variables were calculated from the first three stimuli/contractions, up to the maximum number of usable contractions: 9 (early), 10 (middle), and 15 (late) for the explosive contractions, and 15 for the MVCs. In the torque and EMG averaging methods, dependent variables were averaged over the best 3 out of the first 10 contractions (9 for early-phase explosive) in each condition. For the torque averaging method, the best 3 explosive contractions were those with the greatest torque just prior to MEP (i.e., at 30, 105, and 180 ms from torque on-set, in early, middle, and late conditions, respectively), whilst the best 3 MVCs were those with the highest average torque over 1 s prior to the MEP. For the EMG averaging method, the best explosive contractions were those with the highest EMG RMS amplitude (averaged of all three muscles) from on-set to the start of the MEP (i.e., 0-23, 0-98, 0-173 ms, in early, middle, and late, respectively), whilst the best three MVCs were those with the greatest EMG RMS in the 1 s period before MEP.

### Statistical analysis

Paired t-tests were used to compare the differences between the test and re-test sessions within each condition for each dependent variable. The relative reliability of TMS responses was assessed via the relative test-retest interclass correlation coefficient (ICC) determined via a two-way mixed-effects model, with the absolute agreement, as suggested for a test-retest design (Koo and Li, 2016). ICCs were interpreted as <0.5 poor, ≥0.5 but <0.75 moderate, ≥0.75 and <0.9 good, and ≥0.9 excellent (Koo and Li, 2016). The coefficient of variation (CV) determined the absolute reliability (degree of fluctuation of repeated measurements within individuals) of dependent variables and was calculated for each participant (100 * test-retest standard deviation (SD) / test-retest mean) before being averaged across participants.

Separate ICCs and CVs were produced for each averaging approach; consecutive running averages and the best three based on both torque and EMG averaging methods. Logarithmic functions (*y* = *a* ln(*x*) + k) were fit to data describing the relationship between ICC or CV and the number of consecutive contractions/stimuli over which the dependent variables were averaged to establish trends in the data and enable extrapolation of the ICC or CV beyond the maximum number of useable contractions. Only statistically significant (p<0.05) logarithmic functions are reported in the results. Logarithmic functions were chosen as they provided excellent fits with significant (p<0.05) relationships (r^2^=0.97-0.75 and SD=0.08) and did not cross zero values of CV (as with, e.g., exponential fits), which are physiologically impossible. Best-fit logarithmic functions were made in SPSS v.26 (IBM Inc., Armonk, NY, USA). For ICCs, a confidence interval of 95% was used for the lower bound (Lb) and upper bound (Ub), and CVs were calculated using MATLAB (R2021a Update 5, MathWorks Inc., Natick, MA, USA). The significance level was set at *p* ≤ 0.05. Data are presented in the results as mean ± SD.

## Results

There was no significant difference (p≥0.168) between the test and retest sessions for any of the dependent variables in each condition (Table 1).

**Table 1.**
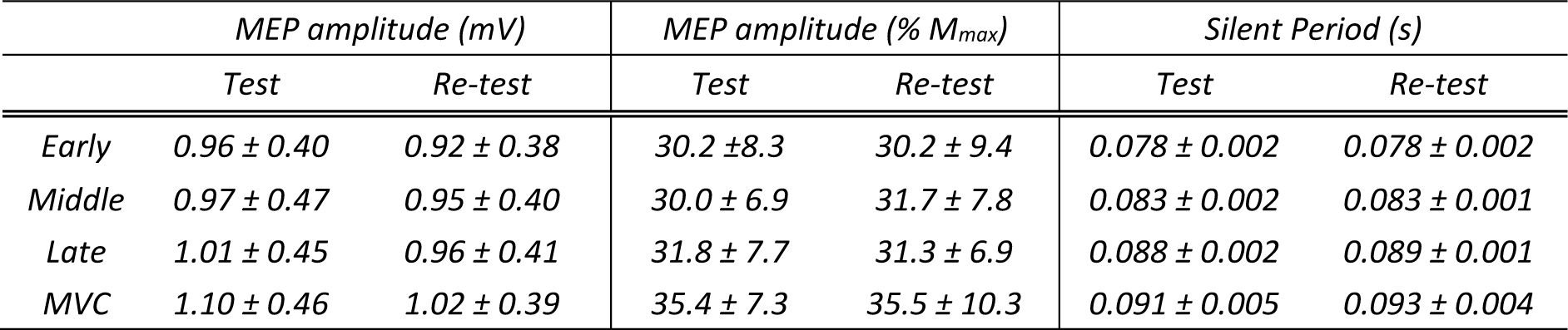
shows the absolute and normalised MEP amplitude, and silent period for each condition in each session (test and re-test). Normalised MEP amplitude is a % of maximal M-wave (M_max_). Conditions are early (45), middle (115), and late (190) phases of explosive contractions, and at the MVC plateau. Data are mean ± SD of the first 10 (9 for early) contractions performed in each condition.

### Reliability as a function of the number of MEPs

The test-retest ICCs for absolute and normalised MEP amplitude generally increased with the number of averaged MEPs (Fig.3 A, B), whilst the test-retest CV for the same variables generally decreased (Fig.3 C, D). These trends indicate increasing test-retest MEP amplitude reliability with increasing numbers of averaged MEPs. The exception to these general trends was the late phase. In this condition, although the CV of normalised MEP amplitude was related to the number of averaged MEPs (Fig 3D), the CV of absolute MEP amplitude, the ICC of absolute MEP amplitude, and the ICC of normalised MEP amplitude were not related to the number of averaged MEPs (Figs 3 A, B, C). These trends, apparent upon visual inspection, were supported by logarithmic fits to the data, which were significant in all cases aside from the late phase cases highlighted in the previous sentence (Table 2).

**Table 2.** Coefficients (a and k), the goodness of fit (R^2^) of the logarithmic functions *y* = *a ln*(*x*) + *k*. Functions were fit to describe the relationship between test-retest ICC or CV and the number of consecutive contractions over which MEP amplitudes were averaged. The ’n of’ is the number of MEP amplitudes averaged to reach ICC≥0.75 or CV≤10%. The * denotes where ’n of’ has been extrapolated beyond the measured data. MEP amplitudes were recorded at the plateau of MVCs, and at three different time points from EMG on-set during explosive contractions: Early (45 ms), middle (115 ms), and Late (190 ms).

| ICC |  |  |  |  |  |  |  |  |  |  |  |
| --- | --- | --- | --- | --- | --- | --- | --- | --- | --- | --- | --- |
| MEP absolute | a | k | R<sup>2</sup> | p | n of | MEP normalised | a | k | R<sup>2</sup> | p | n of |
| Early | 0.116 | 0.704 | 0.96 | 0.001 | 2 | Early | 0.115 | 0.537 | 0.95 | 0.001 | 7 |
| Middle | 0.231 | 0.265 | 0.92 | 0.001 | 9 | Middle | 0.392 | -0.395 | 0.84 | 0.002 | 19* |
| Late | 0.002 | 0.819 | -0.20 | 0.950 | - | Late | -0.093 | 0.660 | 0.13 | 0.220 | - |
| MVC | 0.427 | 0.200 | 0.93 | 0.001 | 4 | MVC | 0.154 | 0.273 | 0.85 | 0.002 | 23* |
| CV |  |  |  |  |  |  |  |  |  |  |  |
| MEP absolute | a | k | R<sup>2</sup> | p | n of | MEP normalised | a | k | R<sup>2</sup> | p | n of |
| Early | -5.17 | 23.38 | 0.90 | 0.001 | 14* | Early | -4.77 | 24.28 | 0.87 | 0.001 | 20* |
| Middle | -7.23 | 29.55 | 0.94 | 0.001 | 15* | Middle | -5.41 | 22.61 | 0.93 | 0.001 | 11* |
| Late | -1.76 | 15.82 | 0.16 | 0.210 | - | Late | -3.04 | 18.71 | 0.95 | 0.001 | 18* |
| MVC | -2.16 | 15.19 | 0.68 | 0.013 | 11 | MVC | -2.90 | 19.04 | 0.79 | 0.005 | 23* |

ICCs were generally higher for absolute MEP amplitude (Fig 3A) than normalised MEP amplitude (Fig 3B). For absolute MEP amplitude, good ICC (ICC>0.75) was found within just three averaged MEPs for all conditions except middle (where it was attained within nine averaged MEPs). For normalised MEP amplitude, good ICC was only achieved in the early condition (attained within seven contractions), whilst other conditions only reached moderate ICCs (ICC>0.50). CV tended to be similar for absolute and normalised MEP amplitudes, reaching 10-15% values for all conditions within seven contractions (Figs 3C, 3D). We were interested in identifying the number of averaged MEPs needed to attain good ICC (ICC>0.75) and CV<10% for all variables and conditions. As is noticeable from Fig 3 A-D, this was not consistently reached using the number of MEPs measured in this study (9-15, depending on the condition). Therefore, the fitted logarithmic functions provided estimates by extrapolating beyond fitted data where necessary (see Table 2 "n of" columns).

In contrast to the general trends for MEP amplitude reliability, the silent period reliability did not vary as a function of number of averaged MEPs (Fig.3E, F). In support, attempted logarithmic fits were all non-significant (p>0.05; fits not shown). Moderate ICCs (0.5≤ICC≤0.75) were generally observed for the silent period in all conditions (Fig 3E). CVs for the silent period (Fig 3F) were lower than for MEP amplitude, particularly during MVC.

### Reliability for best three contractions using EMG or torque methods

Compared to averaging the first 10 consecutive contractions, the ICC tended to be lower (- 0.07±0.01) and CV slightly higher (+1.7±0.2%) for either of the best 3 averaging methods, across all conditions and dependent variables (Table 4). There was no consistency for the torque (best 3) averaging method to be better than the EMG averaging method, nor vice versa, with the higher ICC and lower CV between methods being dependent on condition and variable.

**Table 3.** test-retest ICC, with 95% confidence interval in parenthesis, of dependent variables determined via 3 different averaging methods: average of the first ten consecutive contractions, and average of the best 3 based on those contractions with the highest EMG or torque prior to the MEP. Dependent variables are absolute MEPs, normalised MEPs (normalised to maximal M-wave), and silent period. MEP data were recorded at the plateau of MVCs, and at 3 different time points from EMG on-set during explosive contractions: early (45 ms), middle (115 ms), and late (190 ms)

| *Dependent Variables* | *Averaging Method* |  |  |
| --- | --- | --- | --- |
| *Absolute MEP amplitude* | *First 10* | *Best 3* |  |
|  |  | *EMG* | *Torque* |
| *Early* | 0.95 (0.84-0.98) | 0.93 (0.79-0.98) | 0.88 (0.65-0.96) |
| *Middle* | 0.79 (0.47-0.93) | 0.74 (0.26-0.89) | 0.69 (0.64-0.97) |
| *Late* | 0.85 (0.61-0.95) | 0.83 (0.56-0.94) | 0.84 (0.57-0.94) |
| *MVC* | 0.85 (0.59-0.95) | 0.60 (0.14-0.85) | 0.77 (0.43-0.92) |
| *Normalised MEP amplitude* | *First 10* | *Best 3* |  |
|  |  | *EMG* | *Torque* |
| *Early* | 0.78 (0.39-0.92) | 0.72 (0.34-0.90) | 0.65 (0.21-0.87) |
| *Middle* | 0.55 (0.06-0.83) | 0.37 (-0.1-0.74) | 0.28 (-0.3-0.69) |
| *Late* | 0.49 (-0.05-0.81) | 0.49 (-0.07-0.80) | 0.40 (-0.17-0.76) |
| *MVC* | 0.63 (0.14-0.86) | 0.53 (0.02-0.82) | 0.72 (0.32-0.90) |
| *Silent Period* | *First 10* | *Best 3* |  |
|  |  | *EMG* | *Torque* |
| *Early* | 0.62 (0.14-0.88) | 0.49 (0.15-0.88) | 0.64 (-0.2-0.92) |
| *Middle* | 0.71 (0.28-0.90) | 0.63 (0.21-0.89) | 0.68 (0.16-0.93) |
| *Late* | 0.68 (0.21-0.89) | 0.63 (0.13-0.87) | 0.63 (0.13-0.87) |
| *MVC* | 0.71 (0.23-0.89) | 0.66 (0.20-0.88) | 0.77 (0.42-0.92) |

**Table 4.**
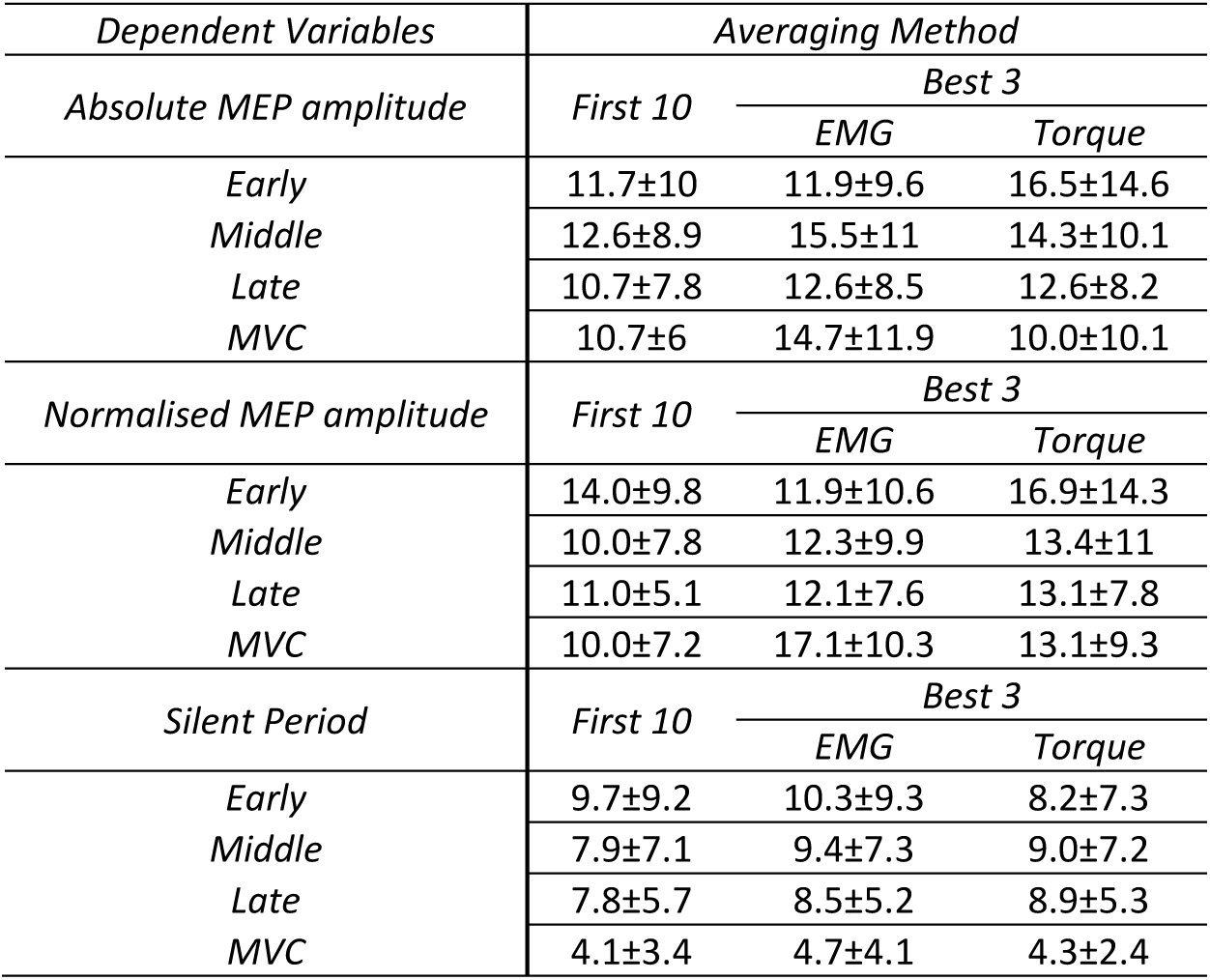
test-retest CV mean ± SD of dependent variables determined via 3 different averaging methods: average of the first ten consecutive contractions, and average of the best 3 based on those contractions with the highest EMG or torque prior to the MEP. Dependent variables are absolute MEPs, normalised MEPs (normalised to maximal M-wave), and silent period. MEP data were recorded at the plateau of MVCs, and at 3 different time points from EMG on-set during explosive contractions: early (45 ms), middle (115 ms), and late (190 ms)

## Discussion

This is the first study to investigate the absolute (ICC) and relative (CV) test-retest reliability of TMS responses (MEP amplitude and silent period duration) recorded at different time points (early, middle, and late) during explosive voluntary contractions, and at the plateau of MVCs. In general, no one condition (explosive early, middle, late, or MVC plateau) produced consistently more reliable (absolute or relative) TMS responses across the different averaging methods than another. In all conditions except the explosive late phase, both absolute and relative reliability of MEP amplitude (absolute and normalised) generally increased with an increase in the number of consecutive stimuli (separate contractions) averaged. In contrast, increasing the number of averaged stimuli did not improve silent period reliability in any condition. This study also found averaging the best 3 out of the first 10 contractions based on EMG amplitude or torque output, produced comparable, albeit slightly poorer, reliability than averaging the same 10 contractions for all TMS responses and conditions.

### Reliability during MVCs

To date, only a handful of studies have investigated the test-retest reliability of MEPs evoked at the MVC plateau, and these have all been based on measures averaged across 3-5 MVCs (Kamen, 2004; Mileva, Sumners and Bowtell, 2012; Souron *et al*., 2016). For comparison within this section, we will focus on our findings within this range of 3-5 contractions, with the influence of higher numbers of contractions being discussed later.

Within 3-5 MVCs, we observed moderate ICC (0.50-0.75) for absolute and normalised MEP amplitudes, which is within the range (0.47-0.79) of what others have observed (Kamen, 2004; Sidhu, Bentley and Carroll, 2009; Mileva, Sumners and Bowtell, 2012; Souron *et al*., 2016; Table 5 in supplementary material). We also observed moderate ICC for silent periods at MVC plateau (0.66-0.70) which was comparable to what has previously been reported for the rectus femoris (0.70; Mileva, Sumners and Bowtell, 2012; 0.61-70; Di Virgilio *et al*., 2022) and lower than reported for the tibialis anterior (0.93-0.95; Souron *et al*., 2016; Table 5 in supplementary material). For test-retest CV, within 3-5 MVCs, we observed 11-16% values for absolute and normalised MEP amplitude and below 5% for the silent period. One previous study has reported test-retest CV for TMS responses at MVC plateau in the tibialis anterior (Souron *et al*., 2016) and reported slightly lower CVs for MEP amplitude (10-11%) and a slightly higher CV for silent period (9%), compared to our results for the quadriceps. A more detailed comparison of our results and those of previous studies is difficult because of the disparity in methods for factors such as the type of ICC used, muscles involved, TMS intensity used, and method of calculating silent period. Nevertheless, based on our results and those of previous studies, averaging data across only 3-5 MVCs should provide moderate ICCs and CVs<16% for MEP amplitudes and silent periods.

Independently of the number of averaged MEPs, absolute MEP amplitude (V) at MVC plateau showed consistently better ICC values than normalised MEP amplitude (% M_max_), and this pattern was consistent for other conditions also. In contrast, a previous study of the tibialis anterior (Souron *et al*., 2016) found ICC for MEP amplitude at the MVC plateau to be similar between absolute and normalised data. This difference between our study and Souron et al. (2016) may be due to the different muscle groups used, although there were other methodological differences (e.g., process for determining coil position and intensity), again making a direct comparison difficult. The greater ICC in absolute compared to normalised MEP amplitude that we observed might originate from the ICC calculation itself. ICC scores will be influenced by inter-participant variability, with greater variability likely contributing to greater ICCs (Koo and Li, 2016). Normalising the MEP amplitudes to M_max_ will control for some of the factors that cause inter-participant variability, such as muscle size and adiposity (Besomi *et al*., 2020), which in turn might reduce ICCs. In contrast to ICC, CV measurements are not influenced by inter-participant variability which may explain why CVs were generally similar for absolute and normalised MEP amplitudes.

### Reliability during explosive contractions

To our knowledge, no prior study has investigated the reliability of TMS responses during explosive contractions. We had suspected the reliability of TMS responses to be lowest in early-phase explosive contractions and highest at the MVC plateau, as this is commonly observed for torque and EMG amplitudes in these different conditions (Folland, Buckthorpe and Hannah, 2014; Tillin and Folland, 2014). However, our findings indicated the test-retest reliability of TMS responses was similar for the different conditions, an observation that appeared unrelated to the number of contractions averaged. Just prior to the onset of rapid contractions, in a period of time that overlaps with the early phase of EMG measurements in the current study, there is an increase in subthreshold cortical excitability and increased probability in evoking a detectable MEP (Mackinnon and Rothwell, 2000). Early subthreshold cortical excitability changes consistently during explosive contractions, but the probability of conversion to detectable voluntary motor output in surface EMG is less reliable (Folland, Buckthorpe and Hannah, 2014; Tillin and Folland, 2014). At later phases in the contraction, corticospinal excitability and voluntary motor output variability align more closely, explaining why early phase EMG and torque measurements are less reliable compared to MEP amplitudes, which remain consistent across phases.

The only exception to there being no clear differences in reliability between conditions was the observation that CV for the silent period was generally highest for the early phase and lowest for the MVC plateau (Fig. 2F). This may be associated with the lower overall length of the silent period in the early phase compared to the MVC plateau (Table 1). Given the CV is a measure of variability relative to the mean, a condition with a lower mean silent period (as is the case for the explosive early phase), will be susceptible to higher CV values where variability is constant. The shorter silent period in the early phase may be due to neural drive in this phase being predominantly feedforward (Škarabot *et al*., 2021) and less affected by afferent feedback or intracortical circuitry causing inhibition.

### Number of averaged MEPs

For absolute and normalised MEPs, ICCs generally increased, and CVs decreased, with an increasing number of contractions, in all conditions except the late phase. This is in agreement with studies on MEP amplitudes recorded at rest or at submaximal force, which also observed improved reliability by increasing the number of averaged stimuli (Goldsworthy, Hordacre and Ridding, 2016; Biabani et al., 2018; Brownstein et al., 2018). Our results show that the conditions in our study, except for the late-explosive phase, require 11-15 contractions (absolute MEP amplitude) and 18-23 contractions (normalised MEP amplitude) to ensure both good ICCs (≥0.75) and CVs≤10%. Further, although caution is required when extrapolating the logarithmic functions reported in this study well beyond the number of contractions measured, the shape of the logarithmic functions indicates little benefit to reliability will be gained by sampling beyond the abovementioned ranges. Variability in the MEP responses in our study are likely affected by the TMS intensity we used (140% AMT), which evokes MEPs in the quadriceps half-way up the impulse-response relationship for the quadriceps (Temesi, Ly and Millet, 2017). As this is the steepest section of the impulse-response relationship, small changes in corticospinal excitability can greatly affect MEP amplitude (Temesi, Ly and Millet, 2017). Combined with the intrinsic fluctuations in ongoing oscillatory activity within the cortical area (Hordacre et al., 2017), this likely explains the inherent variability in MEP amplitudes, which seems to limit test-retest reliability, even with consecutive averaging. Nevertheless, based on our results, we recommend future investigators consider collecting and averaging MEP amplitudes across more than the standard 3-5 MVCs (or explosive contractions) used in previous studies, if this can be managed within their protocol, with an optimal range likely falls between 10-20.

Unlike the other conditions, MEP amplitude reliability in the late-explosive phase (190 ms) was not improved by increasing the number of averaged contractions. This implies that variability in MEP amplitude during the late phase of explosive contraction remains constant and within a narrow range from contraction to contraction, unlike during the earlier phases. We can only speculate about the physiological factors explaining this observation. However, potentially afferent feedback, which will be greater in the later than earlier phases of explosive contraction (Škarabot *et al*., 2021), inhibits the neural drive to muscle and so maybe limits the range of possible MEP amplitudes in the late phase of explosive contraction.

In the silent period, ICC and CV remained constant despite increases in the number of averaged stimuli. Thus, variability in the silent period appears to be consistent and remain within a narrow range of possibilities from contraction to contraction. Factors that are likely contributing to the variability in silent period include the intensity of stimulation – with lower intensities resulting in higher variability (Damron *et al*., 2008) – and differences in motor threshold compared to MEP as changes in response curve reduce variability (Wassermann *et al*., 1993). Our results, obtained at 140% AMT, show that collecting data or more than 3 explosive contractions or MVCs will nt provide meaningful improvements in the reliability of silent period measurament.

In studies on explosive contractions, RFD is often averaged from the best 3 out of as many as 10 or more contractions (Maffiuletti *et al*., 2016). Thus, if intending to relate TMS responses measured during explosive contractions to RFD, it may be pertinent to also average the TMS responses in the best 3 contractions. Of the two methods we used for determining the best 3 (out of the first 10) contractions (torque and EMG methods), neither showed consistently better reliability than the other across all conditions and variables. However, both methods for all conditions provided comparable ICCs and CVs to the average of the first 10 contractions (Tables 3 and 4). It therefore appears that selecting the best 3 contractions to average MEP amplitudes minimises the influence of MEP variability to a similar extent as increasing the number of averaged stimuli over consecutive contractions from 3 to 10. Thus, averaging the best three contractions may be a viable option for investigators in the future, particularly if looking to relate MEP amplitude measurements to RTD.

### Conclusion

Regardless of the averaging method or reliability metric (ICC and CV), no specific condition produced consistently more reliable MEP amplitudes (absolute or normalised), and silent periods than another. Generally, increasing the number of averaged consecutive contractions increased the reliability for MEPs amplitude (absolute and normalised) in all conditions except for late explosive phase, but did not improve the reliability of silent period duration. Thus, it is advisable to collect more than the typical 3-5 contractions, as used by previous studies involving TMS during MVC, to maximise reliability for MEP amplitude; however, 3-5 contractions will suffice for silent period measurements. Alternatively, investigators could opt to average the best 3 contractions based on those with the highest torque or EMG prior to MEP, which provides comparable reliability to averaging the first 10 contractions.

The authors have no relevant financial or non-financial interests to disclose.

## Abbreviations

AMT: Active motor threshold
CV: coefficient of variance
EMG: Electromyography
ICC: Inter-class correlation
MEP: Motor evoked potential
MVC: Maximal voluntary contraction
MU: Motor unit
RF: Rectus femoris
RMS: Root mean squared
RTD: Rate of torque development
SD: Standard deviation
SP: Silent period
TMS: Transcranial magnetic stimulation
VM: Vastus medialis
VL: Vastus lateralis

