## Supplementary Table for "Test-retest reliability of TMS motor evoked responses and silent periods during explosive voluntary isometric contractions"

*Table 1 Between-session ICC for MEP measures evoked and averaged across 3-5 MVCs from current and prior studies.*

| Study | Muscle/s | Number of MEPs | Between-session ICC | | | |
|---|---|---|---|---|---|---|
| | | | ICC Type | Absolute Amplitude | Normalised Amplitude | Silent Period |
| Current study | Average of VM, VL, RF | 3 to 15 | 3,1 mixed | 0.62 [3][a]<br>0.74 [4]<br>0.75 [5] | 0.41 [3]<br>0.52 [4]<br>0.55 [5] | 0.68 [3]<br>0.70 [4]<br>0.66 [5] |
| Kamen et al (2004) | BB | 5 | Not stated | 0.68 | - | - |
| Sidhu et al (2009) | RF | 4 | 2,1 | - | 0.69 | - |
| Mileva et al (2012) | TA[b] | 3 | 1,k | 0.79 | - | 0.93 |
| Souron et al (2016) | TA | 3 | Not stated | 0.47 | 0.52 | 0.95 |
| Di Virgilio et al (2022) | RF | 3 | 2,1 | - | - | 0.7 (general population)<br>0.61 (soccer player) |

***ICC for Normalised MEP amplitude was also reported in Malcolm et al. 2021, but the reported value (0.59) seems to be averaged across multiple contraction intensities (not limited to MVC).***

***Key:*** *[a] Values in square brackets for the current study represent the number of averaged MEPs used to obtain the reported ICCs. [b] study also includes MEPs from Soleus. It is not included here as these were obtained whilst acting as an antagonist.*
***Abbreviations:*** *BB, Biceps brachii; TA, Tibialis anterior, RF, Rectus femoris; VM, Vastus medialis; VL, Vastus lateralis.*
